# The Calcineurin-FoxO-MuRF1 Signaling Pathway Regulates Myofibril Integrity in Cardiomyocytes

**DOI:** 10.1101/130831

**Authors:** Hirohito Shimizu, Adam Langenbacher, Jie Huang, Kevin Wang, Georg W. Otto, Robert Geisler, Yibin Wang, Jau-Nian Chen

## Abstract

Altered Ca^2+^ handling is often present in diseased hearts undergoing structural remodeling and functional deterioration. The influences of Ca^2+^ signaling on cardiac function have been examined extensively, but whether Ca^2+^ directly regulates sarcomere structure has remained elusive. Using a mutant zebrafish model lacking *NCX1* activity in the heart, we explored the impacts of impaired Ca^2+^ homeostasis on myofibril integrity. Gene expression profiling analysis revealed that the E3 ubiquitin ligase MuRF1 is upregulated in *ncx1-*deficient hearts. Intriguingly, knocking down MuRF1 activity or inhibiting proteasome activity preserved myofibril integrity in *ncx1* deficient hearts, revealing a MuRF1-mediated proteasome degradation mechanism that is activated in response to abnormal Ca^2+^ homeostasis. Furthermore, we detected an accumulation of the MuRF1 regulator FoxO in the nuclei of *ncx1*-deficient cardiomyocytes. Overexpression of FoxO in wild type cardiomyocytes induced MuRF1 expression and caused myofibril disarray, whereas inhibiting Calcineurin activity attenuated FoxO-mediated MuRF1 expression and protected sarcomeres from degradation in *ncx1*-deficient hearts. Together, our findings reveal a novel mechanism by which Ca^2+^ overload disrupts the myofibril integrity in heart muscle cells by activating a Calcineurin-FoxO-MuRF1-proteosome signaling pathway.

## Introduction

The establishment and maintenance of rhythmic cardiac contractions require tightly regulated Ca^2+^ signaling and intact contractile machinery. In the heart, a small amount of Ca^2+^ enters cardiomyocytes upon stimulation by an action potential. This Ca^2+^ influx induces the release of a larger amount of Ca^2+^ from the sarcoplasmic reticulum (SR) resulting in an abrupt increase in cytosolic Ca^2+^ levels and muscle contraction. The re-sequestering of Ca^2+^ to the SR by SERCA2 and extrusion of Ca^2+^ from the cell by NCX1 allows the muscle to relax (Bers, 2002). Abnormal Ca^2+^ handling has been associated with cardiac diseases including heart failure and arrhythmia in humans and animal models (Luo and Anderson, 2013) and defective myofibril structures are also often observed in diseased hearts (Lopes and Elliott, 2014). However, whether or not there is a causal relationship between abnormal Ca^2+^ handling and myofibril disarray in diseased myocytes has not yet been established.

The RING finger protein MuRF1 (also known as TRIM63) is a muscle-specific E3 ubiquitin protein ligase involved in the regulation of muscle turnover in normal physiology and under pathological conditions. MuRF1 acts on several sarcomeric target proteins, tagging them with polyubiquitin chains for proteasome-dependent degradation (Kedar et al., 2004; Clarke et al., 2007; Cohen et al., 2009; Mearini et al., 2010). Through this mechanism, MuRF1 regulates normal sarcomere protein turnover and removes misfolded and/or damaged proteins in skeletal and cardiac muscles (Lyon et al., 2013; Pagan et al., 2013; Willis et al., 2014). MuRF1 expression is elevated under muscle catabolic conditions and overexpression of MuRF1 in the heart results in a thin ventricular wall and a rapid transition to heart failure upon transaortic constriction, suggesting that MuRF1 is a major player in muscle catabolic processes (Bodine et al., 2001; Labeit et al., 2010; Baehr et al., 2011; Files et al., 2012; Gomes et al., 2012; Bodine and Baehr, 2014). Conversely, knockout of MuRF1 promotes resistance to muscle atrophy and an exaggerated hypertrophic response to pressure overload (Willis et al., 2007; Willis et al., 2009a; Willis et al., 2009b). In humans, patients with specific MuRF1 gene variants develop hypertrophic cardiomyopathy at a younger age (Chen et al., 2012; Su et al., 2014), revealing a pathological role for MuRF1 in the progression of cardiac diseases.

In skeletal muscles, the Forkhead box O (FoxO) transcription factors serves as a nodal point controlling muscle degradation via regulating MuRF1 expression. Under catabolic conditions, the PI3K-Akt pathway is suppressed and hypophosphorylated FoxO translocates into the nucleus causing MuRF1 induction and muscle atrophy (Lecker et al., 2004; Waddell et al., 2008). Conversely, upon IGF stimulation, AKT is activated to phosphorylate and sequester FoxO in the cytoplasm, resulting in the repression of MuRF1 and an increase in myocyte mass (Sacheck et al., 2004; Stitt et al., 2004). Similarly, an AKT-FoxO-mediated suppression of MuRF1 expression in response to insulin has been noted in cardiac muscles (Skurk et al., 2005; Paula-Gomes et al., 2013).

In this study, we used the zebrafish *tremblor/ncx1h* mutant to explore the regulatory relationship between Ca^2+^ homeostasis and the maintenance of cardiac muscle integrity. We have previously shown that *NCX1h* (also known as *slc8a1a*) encodes a cardiac specific sodium-calcium exchanger 1 (NCX1) in zebrafish and that the *tremblor* mutant lacks functional NCX1h (Langenbacher et al., 2005). NCX1 is a primary Ca^2+^ efflux mechanism in cardiomyocytes (Ottolia et al., 2013), and consistent with this important role in Ca^2+^ homeostasis, cyclic Ca^2+^ transients are abolished in *tremblor/ncx1h* cardiomyocytes resulting in fibrillation-like chaotic cardiac contractions (Ebert et al., 2005; Langenbacher et al., 2005; Shimizu et al., 2015). Like NCX1-/- mice, *tremblor/ncx1h* zebrafish hearts also develop severe myofibril disarray (Koushik et al., 2001; Wakimoto et al., 2003; Ebert et al., 2005), suggesting that a conserved molecular link exists between aberrant Ca^2+^ handling and myofibril disarray. From a microarray analysis, we found that the expression of MuRF1 is significantly upregulated in *ncx1h*-deficient hearts. This MuRF1 upregulation was responsible for the myofibril disarray in *ncx1h*-deficient hearts, and normal cardiac myofibrils could be restored by genetic and pharmacological manipulation of MuRF1 or proteasome activity. We also found that elevated intracellular Ca^2+^ levels enhanced MuRF1 expression via activation of Calcineurin signaling, which dephosphorylates the MuRF1 transcriptional regulator FoxO, leading to its nuclear translocation. Our findings reveal a novel signaling pathway in which Ca^2+^ homeostasis modulates the integrity of cardiac muscle structure via MuRF1 regulation.

## Results and Discussion

### NCX1 is required for the maintenance of myofibril integrity in cardiomyocytes

Zebrafish *ncx1h* mutant embryos lack functional NCX1 in myocardial cells resulting in aberrant Ca^2+^ homeostasis and a fibrillating heart (Ebert et al., 2005; Langenbacher et al., 2005; Shimizu et al., 2015). Similar to the myofibril phenotype observed in NCX1-/- mice, sarcomeres in zebrafish *ncx1h* mutant cardiomyocytes are damaged (Koushik et al., 1999; Wakimoto et al., 2003; Ebert et al., 2005). To investigate whether NCX1 activity affects the assembly or the maintenance of sarcomeres in myocardial cells, we examined the distribution of α-actinin protein. In striated muscles, α-actinin is localized to the Z-line and is a good marker for assessing sarcomere structure. We found that α-actinin is organized into a periodic banding pattern in both wild type and *ncx1h* mutant cardiomyocytes at 30 hpf (Fig. 1A), suggesting that sarcomere assembly is initiated properly in the absence of NCX1 activity. Interestingly, the sarcomeres degenerate in *ncx1h* mutant cardiomyocytes a day later resulting in sporadic distribution of α-actinin (Fig.1A). Zebrafish myocardial cells of the outer curvature normally assume an elongated, flat shape by two days of development (Auman et al., 2007; Cavanaugh et al., 2015). However, *ncx1h* mutant cardiomyocytes fail to elongate (Fig.1B) and both atrial and ventricular chambers become dysmorphic (Fig. 1C), indicating a requirement for NCX1 activity in the maintenance of myofibril integrity and cardiac chamber morphology.

**Figure 1.**
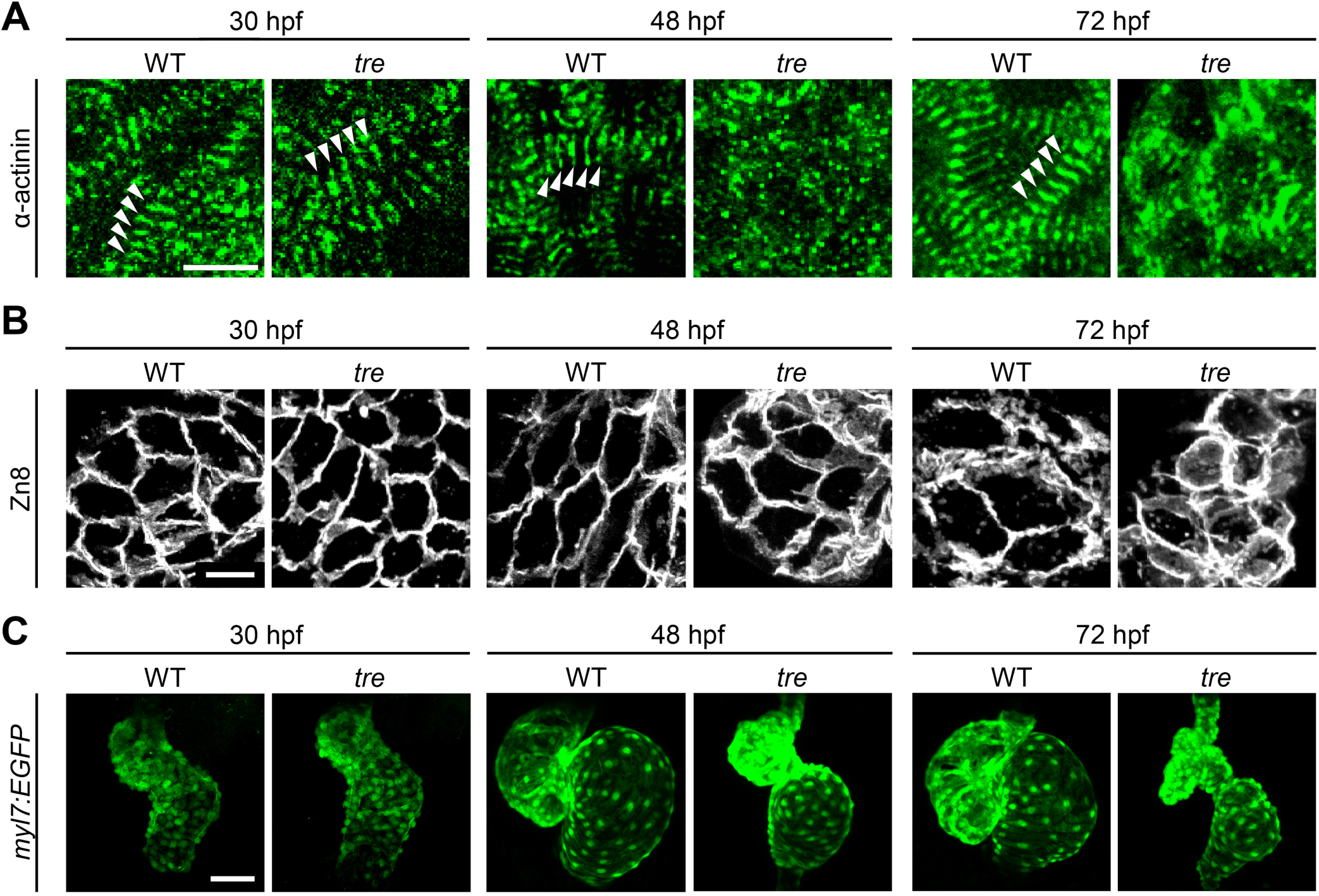
Disorganized myofibril structure in *ncx1* cardiomyocytes. Wild type (WT) and *ncx1h* (*tre*) mutant hearts at 30, 48 and 72 hpf. (A) Zebrafish hearts stained for α-actinin to visualize Z-lines. At 30 hpf, periodic α-actitin staining was observed in wild type and *ncx1h* hearts (arrowheads). By 48 hpf, sarcomeres are disassembled in *ncx1h* hearts. Scale bar, 10 μm. (B) The cell shape of cardiomyocytes was visualized by Zn8 staining. Scale bar, 10 μm. (C) Embryonic fish hearts were visualized by GFP expression in *myl7:EGFP* transgenic background. Note that *ncx1h* hearts become dysmorphic after two days of development. Scale bar, 50 μm.

### Elevated MuRF1 expression in *ncx1h-*deficient hearts

To explore molecular pathways by which NCX1 influences myofibril integrity, we isolated hearts from wild type and *ncx1h* mutant embryos and compared their gene expression profiles. We found that the expression of Muscle Ring-finger protein-1 (MuRF1, also known as TRIM63) is significantly elevated in *ncx1h* mutant hearts. There are two highly homologous MuRF1 genes in zebrafish (MuRF1a/trim63a and MuRF1b/trim63b) (Macqueen et al., 2014). Phylogenetic analysis showed that zebrafish MuRF1a and MuRF1b cluster with other vertebrate MuRF1 genes (Fig. S1A). Both genes span a single exon encoding peptides highly homologous to each other and to their mammalian orthologs (Fig. S1B) (Postlethwait, 2007) and are expressed in striated muscles (Fig.S1C) (Willis and Patterson, 2006). In situ hybridization and quantitative RT-PCR analyses further confirmed that both MuRF1a and 1b are upregulated in *ncx1* mutant hearts (Fig. 2).

**Figure 2.**
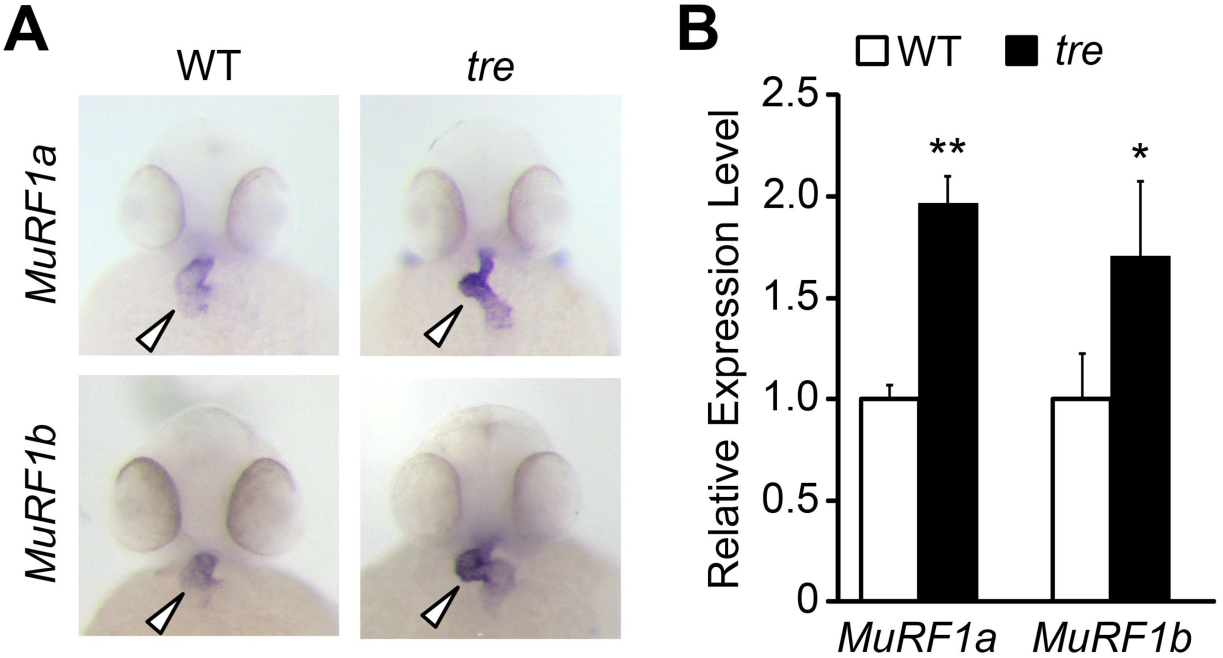
Upregulation of *MuRF1* in *ncx1h* deficient hearts. (A) In situ hybridization analysis shows a significant increase of *MuRF1a* and *MuRF1b* expression in *ncx1* hearts. Arrowheads point to the heart. (B) Quantitative RT-PCR analysis shows an upregulation of MuRF1a and MuRF1b in *ncx1* hearts.

### MuRF1 regulates myofibril integrity in cardiomyocytes

Elevated MuRF1 expression is associated with muscle atrophy and can induce the breakdown of myofibrils in cultured cardiomyocytes (Kedar et al., 2004). We thus asked whether MuRF1 overexpression in the heart is sufficient to induce cardiomyopathy. To this end, we generated a transgenic fish, *myl7:MuRF1a-IRES-GFP*, in which the expression of MuRF1 and GFP reporter is driven by the cardiac-specific *myl7* promoter (Fig. 3A). As shown in Fig.3B, MuRF1 expression is upregulated in *myl7:MuRF1a-IRES-GFP* transgenic hearts. Interestingly, α-actinin failed to organize into a banded pattern in cardiomyocytes of *myl7:MuRF1a-IRES-GFP* embryos (Fig.3B), demonstrating that overexpression of MuRF1 leads to sarcomere disassembly in the heart (Fig.3C). Consequently, MuRF1-overexpressing hearts become dilated (Fig.3D) and their cardiac function is compromised. The fractional shortening of *myl7:MuRF1a-IRES-GFP* hearts was reduced by approximately 10% compared to non-transgenic siblings and the heart rate was also reduced by 10% (Fig. 3E). Together, these findings demonstrate that overexpression of MuRF1 is sufficient to disrupt myofibril structure and impair cardiac function *in vivo*.

**Figure 3.**
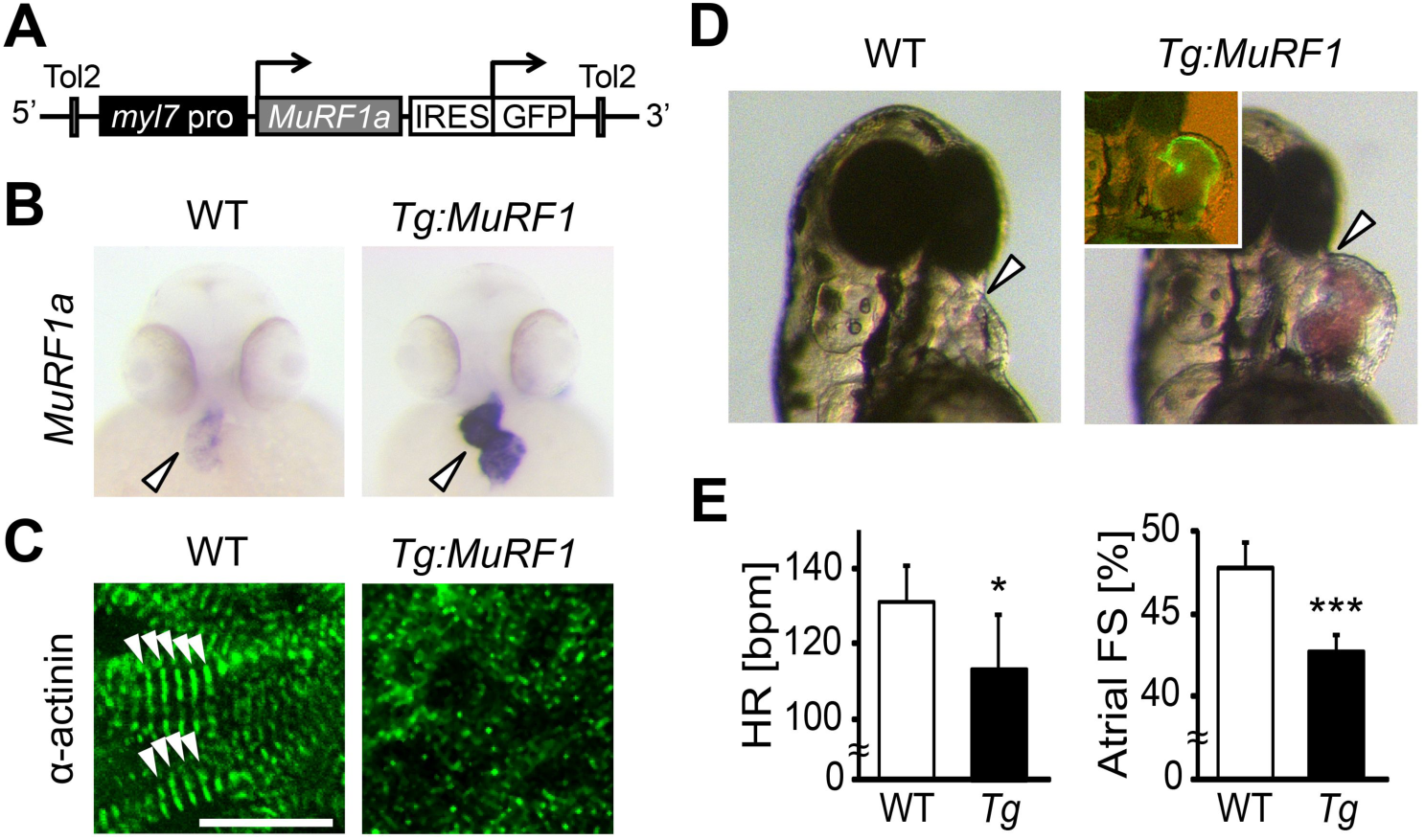
Upregulation of *MuRF1a* leads to myofibril disarray. (A) Schematic representation of the construct that drives cardiac-specific *MuRF1a* expression. (B) *MuRF1a* expression is upregulated in the *myl7:MuRF1a-IRES-GFP* heart (right panel) compared to the wild type heart (left panel). (C) α-actinin staining in wild type (left panel) and transgenic (right panel) cardiomyocytes. Note that sarcomeres are disassembled in *myl7:MuRF1a-IRES-GFP* cardiomyocytes. (D) Live images of wild type and *Tg(myl7:MuRF1a-IRES-GFP)* fish at 72 hpf (left panels). Transgenic hearts are GFP positive and become dilated (inset). (E) Heart rate (HR) and atrial fractional shortening (AFS) in wild type and *myl7:MuRF1a-IRES-GFP* embryos. *, p< 0.05; ***, *p*< 0.001.

### Blocking MuRF1-induced protein degradation preserves myofibril integrity in *ncx1h* mutant hearts

MuRF1’s upregulation upon loss of *ncx1h* activity, along with its established function as a muscle-specific E3 ubiquitin protein ligase that targets sarcomeric proteins for proteasome degradation, make it a good candidate for the cause of the myofibril disarray present in *ncx1h* deficient hearts. If MuRF1 upregulation indeed causes sarcomere disassembly, one would predict that blocking MuRF1 activity or its downstream protein degradation pathway might ameliorate the myofibril defects in *ncx1* mutant hearts. Since both MuRF1a and MuRF1b are upregulated in *ncx1h* deficient hearts, we knocked down these genes simultaneously. We found that ∼80% of *ncx1h/murf1a/murf1b* triple-deficient embryos had intact sarcomeres (n=24), a significant increase compared to *ncx1h* mutant hearts (∼35%, n=21; *p*< 0.001) (Fig. 4). Similarly, treatment with the proteasome inhibitor MG132 suppressed the myofibril disarray caused by NCX1 deficiency. Approximately 72% of MG132-treated *ncx1h* mutants had a banded pattern of α-actinin indicating intact sarcomeres (n=18; *p*< 0.001) (Fig. 4), suggesting that upregulation of MuRF1 induces myofibril degradation via a proteasome-dependent mechanism in *ncx1h*-deficient cardiomyocytes.

**Figure 4.**
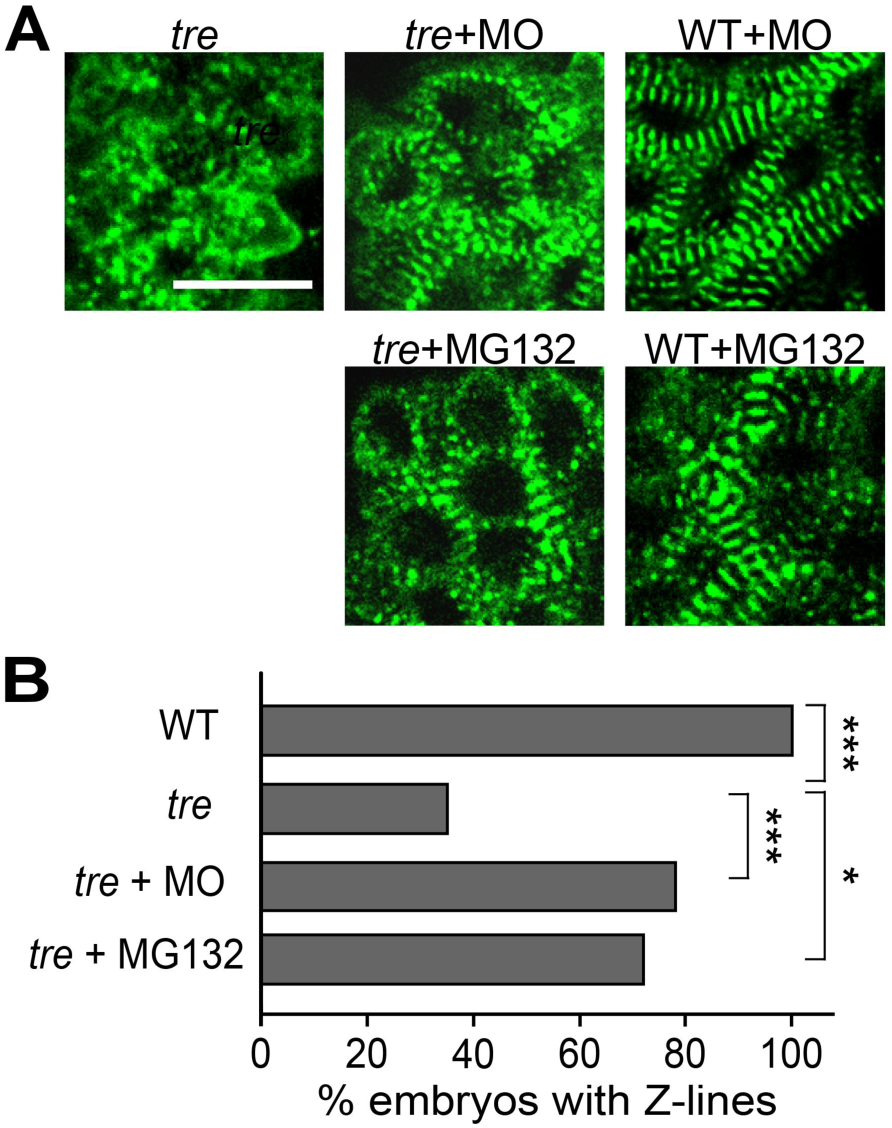
Blocking MuRF1-mediated proteasome degradation preserves myofibril integrity in ncx1 deficient hearts. (A) Z-lines were visualized by α-actinin staining. By 72 hpf, sarcomeres are disassembled in *ncx1h* hearts. MuRF1a/MuRF1b knockdown does not affect sarcomere integrity in wild type embryos (WT+MO), but prevents sarcomere degradation in *ncx1h* (*ncx1h*+MO). Similarly, treatment with a proteasome inhibitor, MG132, preserves myofibril integrity in *ncx1h* cardiomyocytes (*ncx1h*+ MG132). Scale bar, 10 μm. (B) Graph shows % of embryos with periodic α-actinin staining. Chi-squared test, *, *p*< 0.05; **, *p*< 0.01; ***, *p*< 0.001.

### Ca^2+^ induces MuRF1a expression

Our study showed that while *ncx1h* mutant hearts never establish normal Ca^2+^ cycles or heartbeats, the initial assembly of sarcomeres proceeds properly. Since the Ca^2+^ handling defects precede the breakdown of sarcomeres, we hypothesized that Ca^2+^ overload induces MuRF1 expression in cardiomyocytes and thereby leads to sarcomere disassembly and cardiomyopathy. To examine this hypothesis, we isolated a 6.9 kb genomic fragment upstream of MuRF1a, MuRF1a (-6906). Transgenic analysis showed that this genomic fragment was sufficient to drive GFP expression in cardiac and skeletal muscles (Fig. 5A and B), a pattern resembling the endogenous MuRF1 expression pattern (Fig. S1C), indicating that critical regulatory elements are present in this genomic fragment. We then created a MuRF1a (-6906) Luciferase reporter construct to test whether this MuRF1 upstream regulatory element is responsive to Ca^2+^ signaling. We transfected the MuRF1a (-6906) Luciferase reporter into HEK293T cells and induced Ca^2+^ flux by treatment with the Ca^2+^ ionophore A23187. Interestingly, the luciferase activity was significantly enhanced by A23187 induction (Fig. 5C), demonstrating that MuRF1 transcription is sensitive to intracellular Ca^2+^ levels.

**Figure 5.**
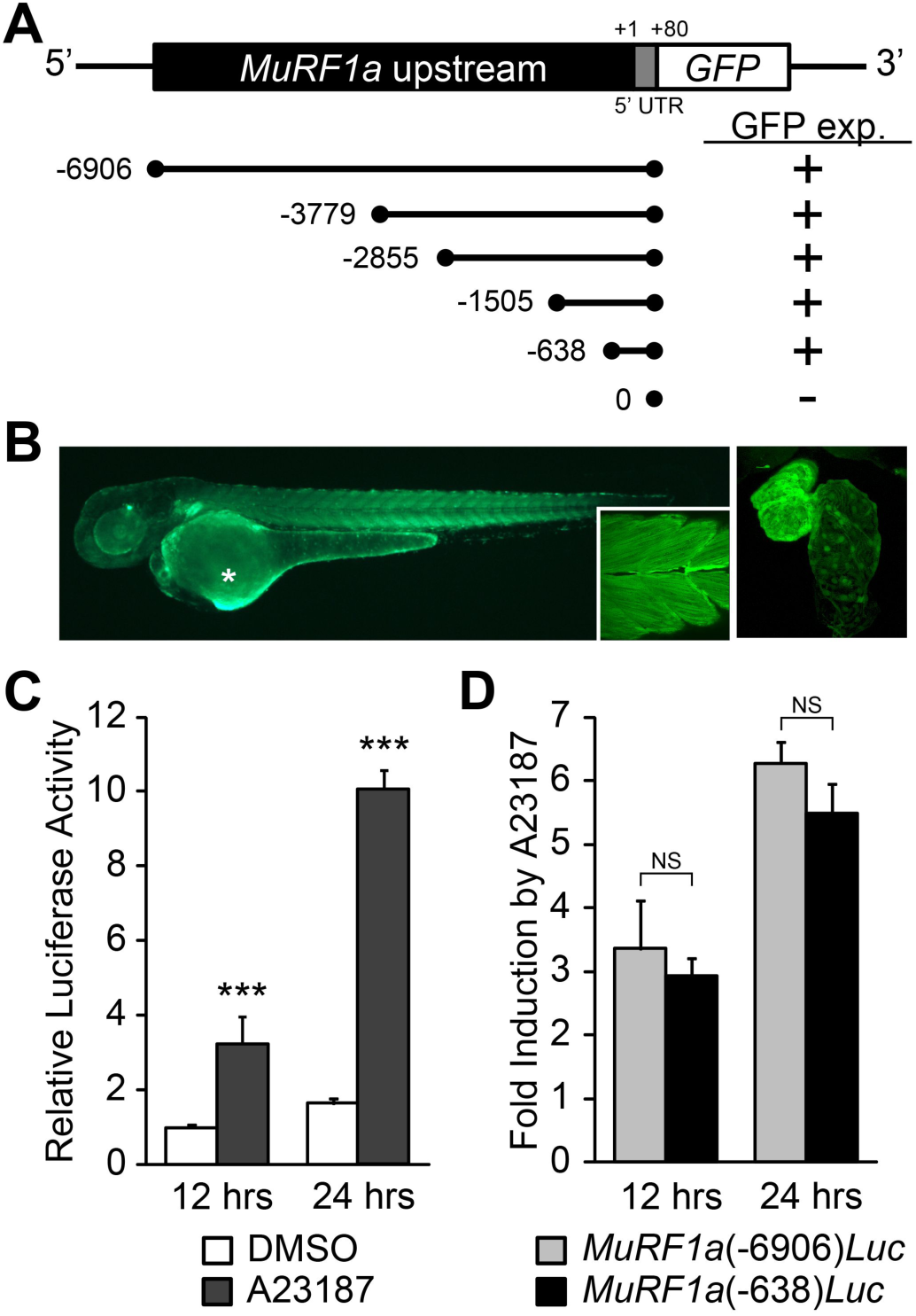
Identification of MuRF1a regulatory element. (A) Schematic representation of MuRF1a reporter constructs. + denotes the presence of GFP expression in the heart and somites. (B) MuRF1a (- 6906)-GFP embryo shows GFP expression in the heart and the somites (inset shows higher magnification image of the somites). The right panel shows a higher magnification image of the heart. The asterisk denotes auto-fluorescence from the yolk. (C) A23187 treatment induces luciferase activity driven by MuRF1a (-6906) promoter. Values represent the fold increase compared to cells treated with DMSO for 12 hours. (D) Comparison of Ca^2+^ responsiveness between MuRF1a (-638) and MuRF1a (-6906) promoters. Values represent A23187 treatment induced fold increase of luciferase activities. ***, *p*< 0.001; NS, not significant.

We next tested the expression patterns of a series of deletion constructs of MuRF1a (-6906) and found that the 638 bp region immediately upstream of the transcription initiation site, MuRF1a (-638), was sufficient to drive reporter gene expression in cardiac and skeletal muscles. MuRF1a (-638)-Luc also displayed enhanced luciferase activity upon A23187 induction at levels comparable to MuRF1a (-6906)-Luc (Fig. 5D), indicating that the 638bp MuRF1a proximal region is sufficient to direct Ca^2+^-mediated MuRF1 transcription.

### Ca^2+^ regulates MuRF1 expression via the Calcineurin-FoxO signaling pathway

Calmodulin-dependent protein kinase II (CaMKII) and the calmodulin-dependent protein phosphatase calcineurin (Cn) are two major transducers of Ca^2+^ signals in cardiomyocytes (Heineke and Molkentin, 2006; Maillet et al., 2013). We asked whether either of these pathways is involved in the regulation of MuRF1 gene expression. We treated MuRF1a (-638)-Luc-transfected HEK293T cells with either KN62, a chemical inhibitor of CaMKII, or FK506, an inhibitor of Cn. KN62 treatment did not have a significant impact on MuRF1a (-638)-driven expression, but FK506 treatment potently attenuated the A23187-induced increase of MuRF1a (-638)-Luc reporter activity (Fig. 6A), suggesting that MuRF1 expression is regulated by a Cn-mediated mechanism. This interpretation is further supported by the observations that Cn overexpression enhances A23187-induced MuRF1 reporter activity and that the A23187-induced MuRF1 expression is blunted by overexpression of a dominant negative form of Cn (DN-Cn) (Fig. 6B) (Kahl and Means, 2004).

**Figure 6.**
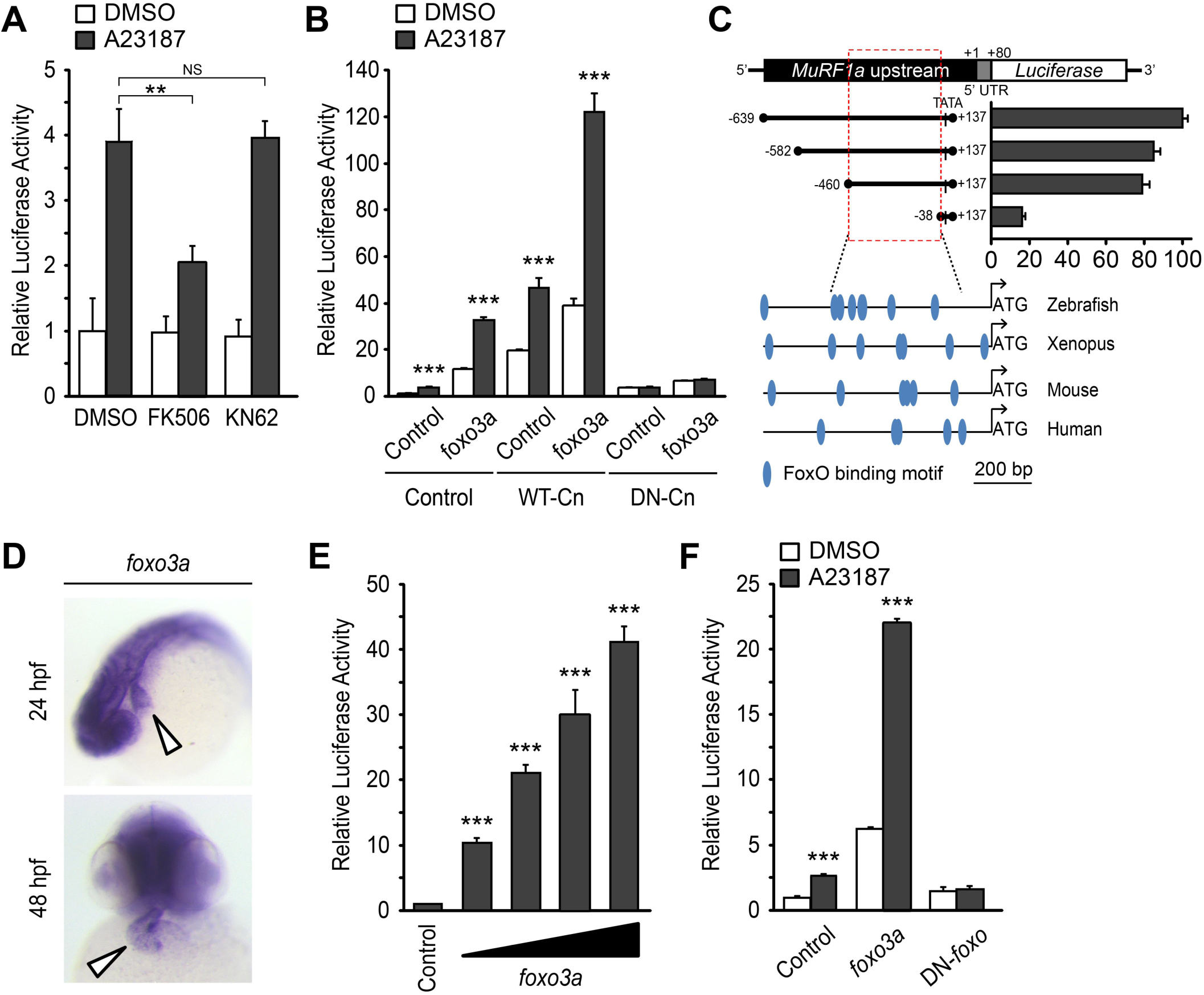
Ca^2+^-Cn-FoxO signaling pathway regulates MuRF1 expression. (A) HEK293T cells were transiently transfected with the MuRF1a (-638) luciferase reporter construct. Cells were incubated with FK506 or KN62 before DMSO or A23187 treatment. (B) Luciferase assay of the MuRF1a (-638) reporter cotransfected with *foxo3a*, wild type calcineurin (WT-Cn) or dominant-negative calcineurin (DN-Cn). (C) Diagrams represent serial deletion of the MuRF1a (-638) reporter. Bar graphs show luciferase activity of each reporter construct relative to that of the empty expression plasmid. Red dotted box indicates the minimal *cis*-regulatory element of MuRF1a. Lower diagrams represent the alignment of zebrafish, *Xenopus*, mouse and human MuRF1 promoters. Blue circles indicate putative FoxO binding sites (D) Whole-mount in situ hybridization detects foxo3a expression in the zebrafish heart. White arrowheads point to the heart. (E) HEK293T cells were transfected with the MuRF1a (-638) luciferase reporter and *foxo3a* expression plasmid. (F) HEK293T cells were transfected with MuRF1a (- 638) reporter plasmid and either wild type or dominant negative foxo3a expression plasmid. Values are expressed relative to the luciferase activity of DMSO treated cells. **, *p*< 0.01; ***, *p*< 0.001; NS, not significant.

We next explored the molecular link by which Cn influences MuRF1 expression. We found multiple putative FoxO binding sites located the minimal regulatory regions of zebrafish MuRF1a/b and within the 1 kb region immediately upstream of the transcription initiation sites of the frog, mouse and human MuRF1 genes (Fig. 6C). Since FoxO is a downstream mediator of Cn signaling (Hudson and Price, 2013) and is involved in the regulation of MuRF1 in skeletal muscles (Stitt et al., 2004; Waddell et al., 2008), we examined the possibility that FoxO mediates Cn’s regulation of MuRF1 expression in cardiomyocytes. There are seven FoxO genes in zebrafish (Wang et al., 2009), all of which are expressed in the developing heart (Fig.6D and Fig. S2). When co-transfected with the MuRF1a(-638)-Luc reporter, all zebrafish FoxO genes tested were capable of enhancing the A23187 induction of MuRF1a(-638) luciferase activity (Fig. S3). FoxO3a promoted strong MuRF1a promoter activity (Fig. S3) and was used for the remainder of our analyses. We found that FoxO3a enhanced MuRF1a(-638)-Luc reporter activity in a dose-dependent manner (Fig. 6E) whereas overexpression of a dominant-negative form of FoxO (DN-foxO), which lacks the transactivation domain but harbors an intact DNA binding domain (Medema et al., 2000; van den Heuvel et al., 2005), abrogated the A23187-induced MuRF1a(-638)-Luc activity (Fig. 6F). We next asked whether FoxO mediates Cn signaling to control MuRF1 expression. We found that cotransfection of Cn and FoxO3a enhances MuRF1a promoter activity (Fig. 6B, E) and that FoxO could no longer induce MuRF1a expression in the presence of a dominant negative form of Cn (Fig. 6B, E), demonstrating that Ca^2+^ influences MuRF1 expression via the Cn-FoxO signaling axis.

### Cn and FoxO regulate MuRF1 expression in the heart

Based on our finding that a Cn-FoxO-MuRF1 regulatory pathway is activated in response to elevated intracellular Ca^2+^ levels in cultured cells, we explored the significance of the Cn-FoxO-MuRF1 pathway in the regulation of myofibril integrity in myocardial cells *in vivo*. The subcellular localization of FoxO is controlled by its phosphorylation status (Huang and Tindall, 2007). We reasoned that the Ca^2+^ extrusion defect in *ncx1h* mutant hearts could activate Cn resulting in the dephosphorylation and nuclear translocation of FoxO. Indeed, while FoxO was primarily sequestered in the cytoplasm of cardiomyocytes in wild type zebrafish hearts, FoxO protein was enriched in the nuclei of *ncx1h* mutant cardiomyocytes (Fig. 7A). This nuclear accumulation of FoxO correlated with the increased MuRF1 expression in *ncx1h* mutant hearts (Fig. 2). In addition, we found that MuRF1 expression could also be induced in the heart by overexpression of FoxO3a or a constitutively active form of FoxO3a in which three phosphorylation sites were replaced by with alanines (CA-FoxO3a: T29A, S236A, S299A) (Fig. 7B) (Brunet et al., 1999). Conversely, pharmacological inhibition of Cn activity by treatment with FK506 or overexpression of DN-FoxO blunted MuRF1 expression in *ncx1h* mutant embryos (Fig. 7B). Finally, we used α-actinin as a proxy to examine whether the correlation between FoxO and MuRF1 expression translates to the preservation of sarcomere structure. We found that overexpression of CA-FoxO3a in wild type embryos resulted in a sporadic α-actinin distribution in cardiomyocytes that resembled the phenotype observed in *ncx1h* mutant hearts whereas overexpression of DN-FoxO restored a periodic α-actinin banding pattern in *ncx1h* mutant hearts (Fig.7C).

**Figure 7.**
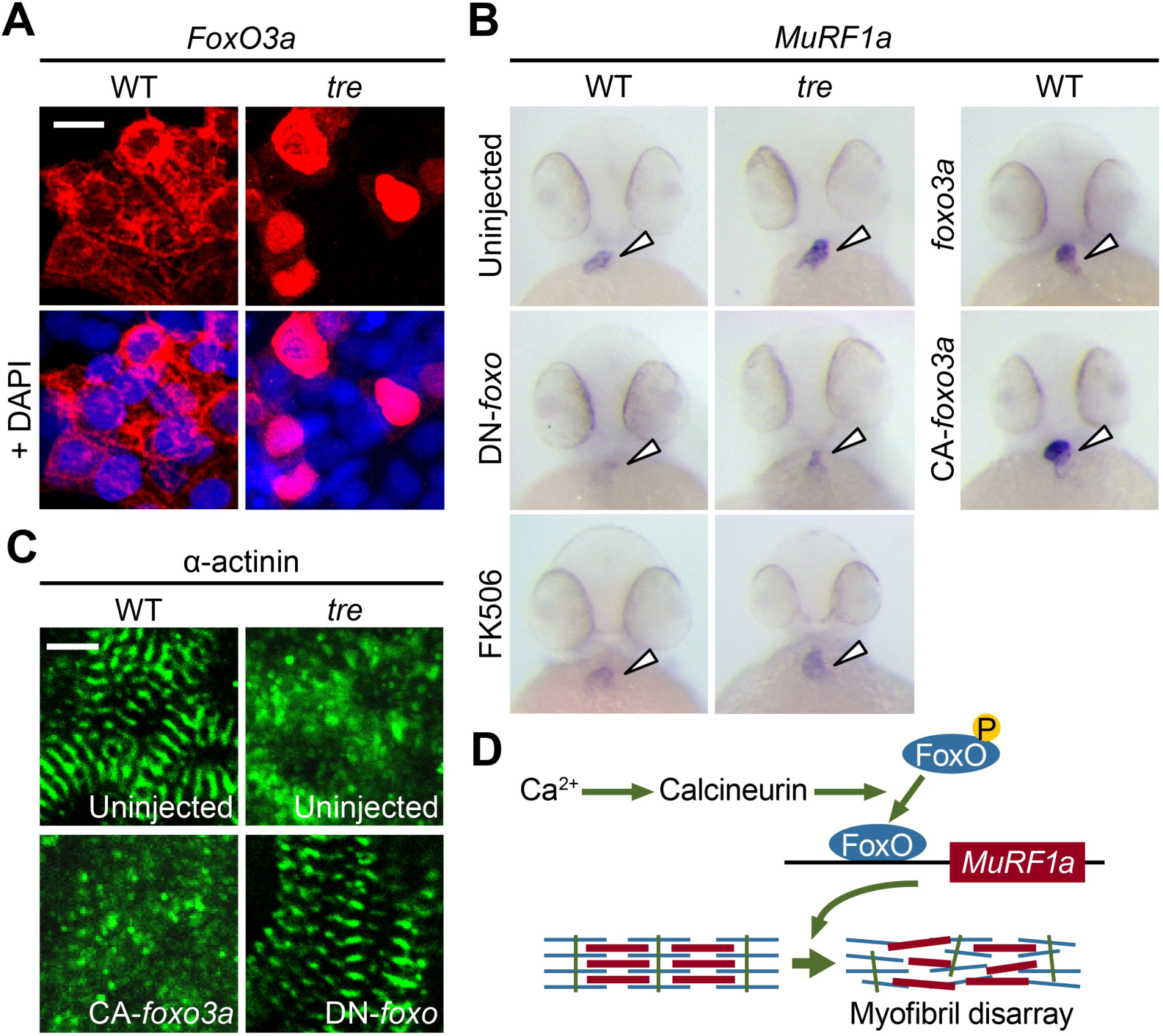
FoxO3a regulates MuRF1 expression in the heart. (A) FoxO3 is predominantly localized in the cytoplasm of wild type cardiomyocytes, but is accumulated in the nuclei of *ncx1h* cardiomyocytes. FoxO3 is pseudo colored in red and nuclei are labeled by DAPI in blue. Scale bar: 10 μm. (B) In situ hybridization showing stronger MuRF1 signals in *ncx1h* mutant and CA-foxo3a injected hearts compared to wild type siblings. MuRF1a expression in *ncx1h* hearts is suppressed by DN-foxo overexpression or FK506 treatment. (C) Immunostaining for the α-actinin in 2 dpf hearts. Intact sarcomeres were detected in control (uninjected) and DN-foxo3a injected *ncx1h* hearts whereas disassembled sarcomeres were observed in *ncx1h* and CA-foxo3a injected hearts. Scale bar: 5 μm. (D) Model for Ca^2+^ overload-induced myofibril disarray. Calcineurin dephosphorylates FoxO leading to FoxO nuclear translocation, MuRF1 expression and sarcomere disassembly.

## Conclusion

Compromised Ca^2+^ homeostasis and damaged cardiac muscles are often observed in deteriorating diseased hearts, but a causative relationship between these outcomes has not previously been demonstrated. In this study, we used the zebrafish *ncx1h* mutant as an animal model to explore the molecular link between Ca^2+^ signaling and myofibril integrity in the heart. We showed that NCX1 activity is dispensable for the initial assembly of sarcomeres, but the maintenance of myofibril structure in myocardial cells requires tightly controlled Ca^2+^ homeostasis and MuRF1 expression.

Our molecular analyses using cultured cells and *in vivo* studies in zebrafish reveal a FoxO-MuRF1 signaling axis that is critically involved in the Ca^2+^-dependent regulation of myofibril integrity in the heart. We propose that under normal physiological conditions where the cytosolic diastolic Ca^2+^ level is low, FoxO is sequestered in the cytoplasm and MuRF1 expression is maintained at a basal level to support the normal turnover of sarcomeric proteins. Under pathological conditions, when diastolic Ca^2+^ is elevated, the activation of Cn dephosphorylates FoxO and allows its nuclear translocation, leading to upregulation of MuRF1 and the degradation of myofibrils (Fig.7D). Interfering with the Cn-FoxO-MuRF1-proteosome pathway by pharmacological or genetic means can protect the sarcomeric integrity of cardiomyocytes suffering from Ca^2+^ dysregulation, and suggest that the FoxO-MuRF1 signaling axis is a central regulator of the Ca^2+^-dependent growth and degradation of striated muscles. The activity of the Cn-FoxO-MuRF1 signaling pathway identified in this study is consistent with the roles of the FoxO-MuRF1 pathway in hypertrophy and atrophy responses in skeletal muscles (Sacheck et al., 2004; Stitt et al., 2004; Waddell et al., 2008) and suggests that FoxO-MuRF1 signaling is critical in the maintenance of tissue homeostasis and in response to pathological stimuli. Furthermore, cardiac-specific overexpression of MuRF1 results in phenotypes resembling those observed in cardiomyopathy; the breakdown of sarcomeres and a dilated heart with reduced heart rate and decreased contractility, suggesting that misregulation of MuRF1 may contribute to pathological progression of cardiovascular diseases. Interestingly, cardiac patients carrying specific MuRF1 gene variants have a poor prognosis (Chen et al., 2012; Su et al., 2014), suggesting that MuRF1 has a conserved role in the regulation of cardiac structure and function from lower vertebrates to humans and raising an intriguing possibility that the Cn-FoxO-MuRF1-proteosome pathway may be an attractive point of therapeutic intervention for cardiomyopathies.

## Materials and Methods

### Zebrafish husbandry, chemical treatment and morpholino knockdown

Zebrafish *tremblor*^*tc318*^ heterozygotes were bred in *Tg(myl7:EGFP)* background and raised as previously described (Westerfield, 2000). Embryos were raised at 28.5 °C and staged as previously described (Kimmel et al., 1995). For Cn or proteasome inhibition, embryos were treated with 10 μM FK506 (Sigma-Aldrich) or 50 μM MG132 (Calbiochem) at 24 hpf. The morpholino-modified antisense oligonucleotides targeting the translation initiation sites of *MuRF1a* and *MuRF1b* (Table S1, Gene Tools) were microinjected at 1- to 2-cell stage (8ng each).

### Zebrafish transgenesis

Transgenic constructs, *myl7*:*MuRF1a*-*IRES*-*EGFP* and *myl7*:*FLAG*-*foxo3a*-*IRES*-*EGFP*, were generated using the Tol2kit (Kwan et al., 2007). Wild type embryos were injected at the 1-cell stage with 10-20 pg of the transgene plasmid and 20 pg of mRNA encoding Tol2 transposase. Embryos with cardiac-specific EGFP expression were raised as founders.

### Microarray and quantitative PCR

Wild type and *tre* mutant hearts were isolated at 48 hpf as previously described (Burns and MacRae, 2006). Total RNA was purified using RNAeasy micro kit (Qiagen). Microarray hybridization was performed in triplicate using the Affymetrix Zebrafish GeneChip containing 15,617 genes. Data were analyzed using scripts written in the statistical programming language R (Team, 2014). Differentially expressed genes were identified using linear models and multiple testing correction implemented in the Limma package (Smyth, 2004). The relative expression levels of *MuRF1a* and *MuRF1b* in the wild type and *tre* hearts were determined by quantitative PCR using the LightCycler 480 System (Roche Applied Science). GAPDH served as the internal control for normalization. Primer sequences used in this study are listed in Table S1.

### In vivo GFP reporter assay

An approximately 7.0-kb genomic fragment upstream of the zebrafish *MuRF1a* gene (ranging from –6906 to +80 bp) was amplified from genomic DNA. A deletion series of *MuRF1a*-EGFP construct was generated using the ERASE-A-BASE system (Promega). For transient expression analysis, each deletion construct was digested with NheI and SalI to release the *MuRF1*a-EGFP reporter and microinjected into 1-cell stage embryos. A minimum of 20 EGFP-positive embryos of each group were examined at 1 and 2 days post fertilization using a Zeiss SV-11 epifluorescence microscope.

### Whole-mount in situ hybridization and immunostaining

Whole mount in situ hybridization and immunostaining were performed as preciously described (Cavanaugh et al., 2015). The antisense RNA probes were synthesized from pCS2+ expression constructs containing a partial genomic fragment (*foxo5a*) or full-length cDNA fragments (*MuRF1a*, *MuRF1b*, *foxo1a*, *foxo1b*, *foxo3a*, *foxo3b*, *foxo4*, and *foxo5b*). Phalloidin (1:50, Sigma-Aldrich) and anti-sarcomeric α-actinin (1:1000, clone EA53, Sigma-Aldrich), α-FLAG (1:100, clone M2, Sigma-Aldrich) and Zn8 (1:100, Developmental Studies Hybridoma Bank) were used for immunostaining. Fluorescence images were acquired using LSM 510 confocal microscope (Zeiss) with a 40x water objective.

### Cardiac imaging and analysis

Videos of *Tg(myl7:MuRF1-IRES-EGFP)* and *Tg(myl7:EGFP)* hearts were taken at 30 frames per second. Cardiac parameters were assessed by line-scan analysis as previously described (Shimizu et al., 2015). Embryos in the *Tg(myl7:gCaMP4)* background was used for Ca^2+^ imaging as previously described (Shimizu et al., 2015).

### Cell-based luciferase assay

HEK293T cells were plated into 96-well plates at a density of 32000 cells per well and transfected with 200 ng of the *MuRF1a* (-6906)- or *MuRF1a* (-638)-luciferase reporter construct, 50 ng of the SV40-*Renilla* luciferase reporter construct and expression vectors (Cn, DN-Cn, foxo3a, CA-foxo3a or DN-foxo3a). Cells were treated with 5μM A23187 (Sigma-Aldrich), 0.5μM FK506 (Sigma-Aldrich) or 0.5μM KN62 (Sigma-Aldrich). Luciferase activities were determined by the Dual-Glo Luciferase Assay System (Promega) in triplicate at least three times, and the activity of firefly luciferase was normalized to that of *Renilla* luciferase for transfection efficiency and cell viability.

### Statistics

Data are presented as the mean ± S.E. *p*-values associated with all comparisons are based on Student’s t-tests (n≥3) unless otherwise stated.

## Acknowledgements

The authors thank members of the Chen Lab for stimulating discussions. This work was supported by grants from the Nakajima Foundation (to HS), the National Institute of Health (HL096980 and HL126051 to JNC and HL108186 to YW), European Commission’s Sixth Framework Programme (ZF-MODELS project to RG) and Seventh Framework Programme (ZF-HEALTH project to RG).

The authors declare no competing financial interests.

---

**Figure S1.**
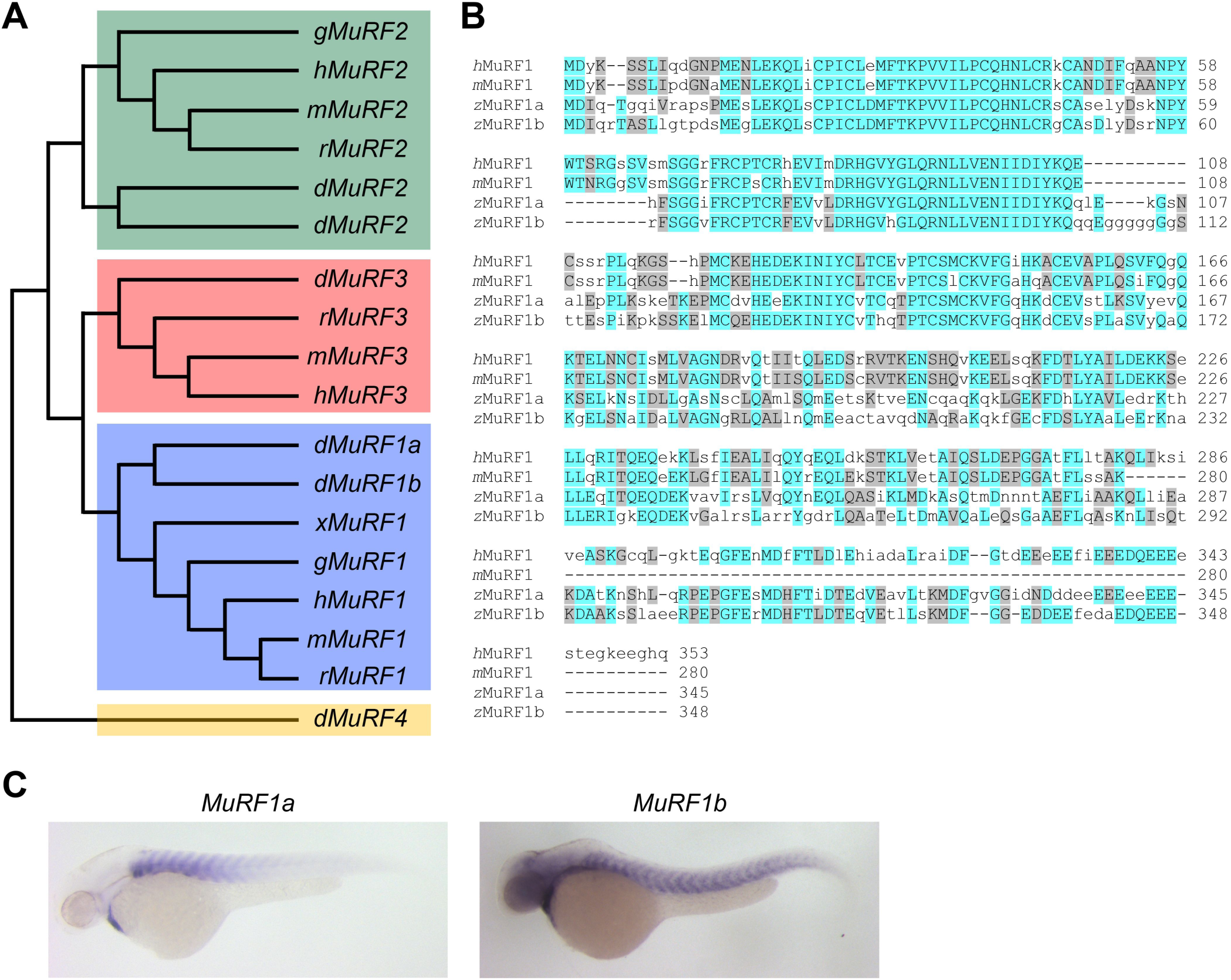
Zebrafish MuRF1 genes. (A) Phylogenetic tree of vertebrate *MuRF1*, *2, 3* and *4* (also known as *trim63*, *55*, *54* and *101*, respectively). The tree was constructed using ClustalX with the neighbor-joining method. Zebrafish (*z*), Human (*h*), mouse (*m*), rat (*r*), chick (*g*), frog (*x*). (B) Alignment of *MuRF1* genes from human, mouse and the zebrafish. Blue boxes highlight identical amino acids and grey boxes indicate similar residues. (C) Whole-mount in situ hybridization demonstrating the expression patterns of MuRF1a and 1b in the zebrafish embryo.

**Figure S2.**
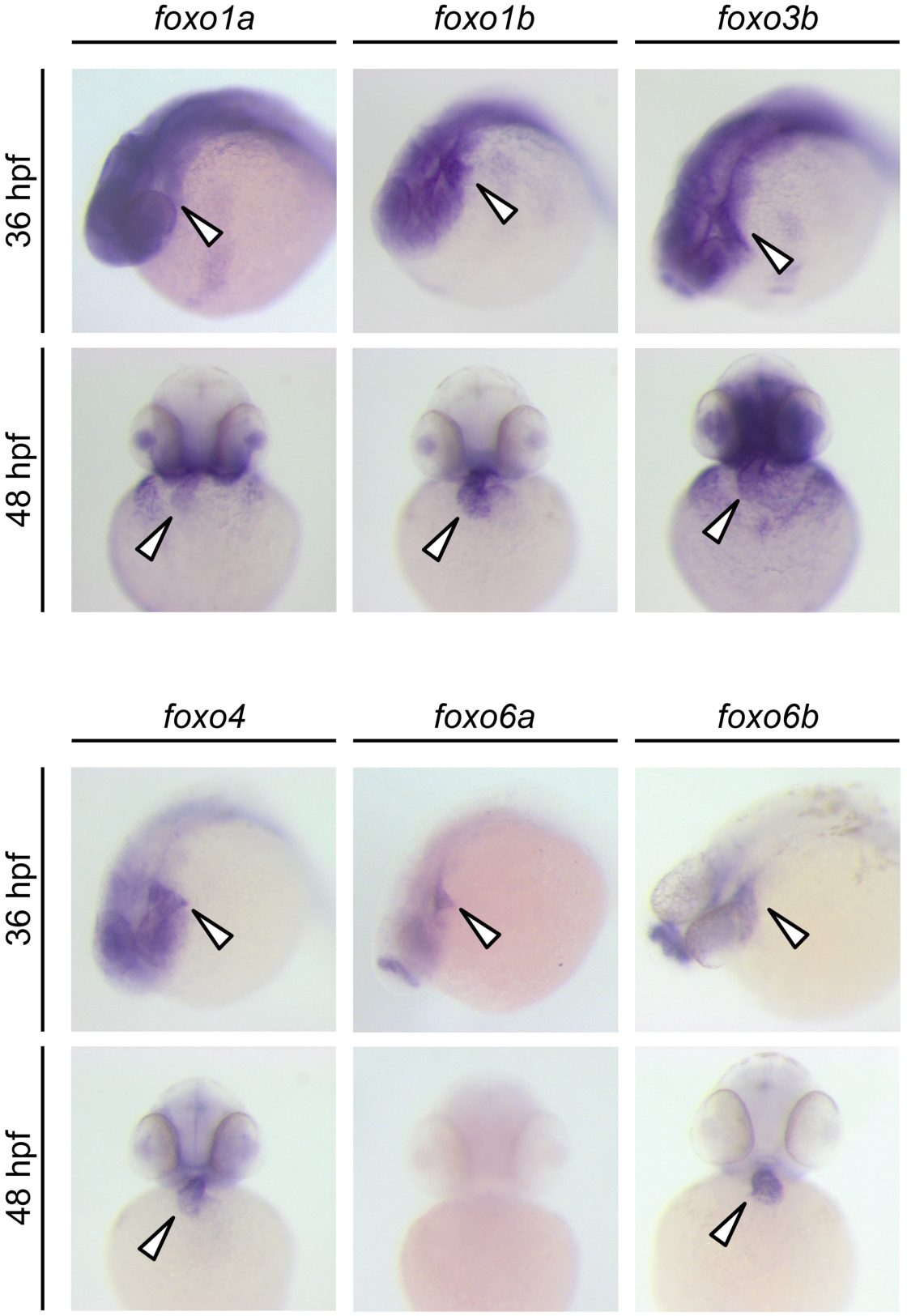
Expression patterns of zebrafish *foxo* genes. Whole-mount in situ hybridization analysis showing *foxo* expression in zebrafish embryos. While *foxo6a* expression is diminished by 48 hpf, all the other *foxo* genes examined (*foxo1a*, *1b*, *3b*, *4*, *6a* and *6b*) are persistently expressed in the heart. Arrowheads point to the heart.

**Figure S3.**
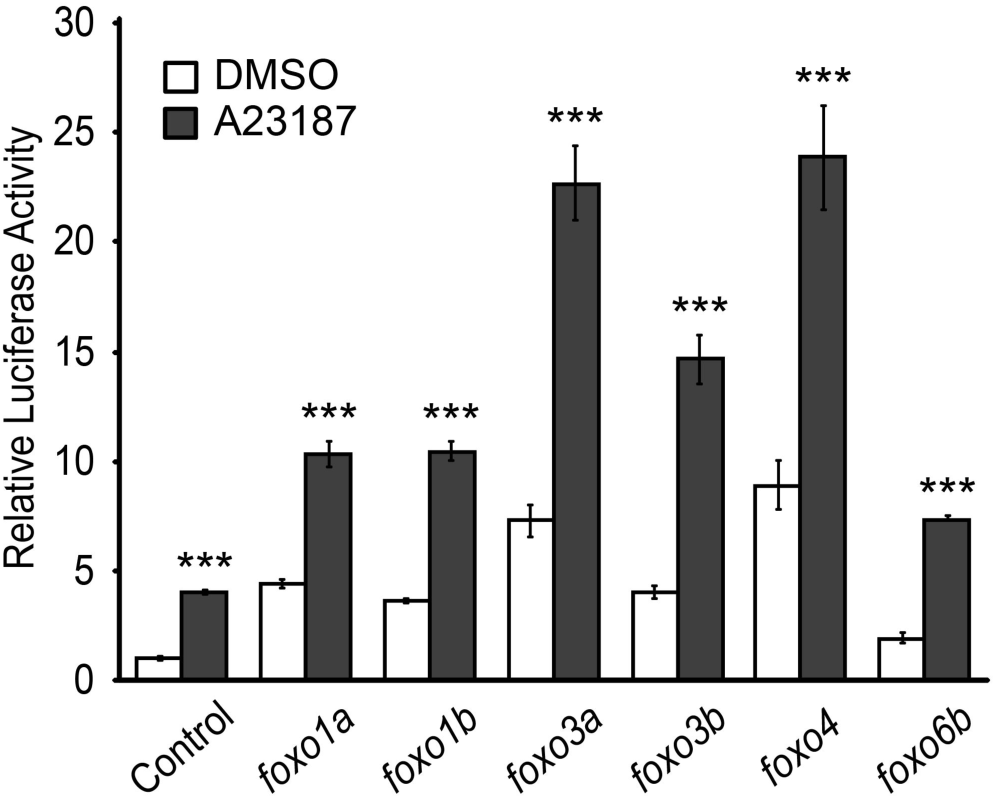
FoxO induces MuRF1 expression. Luciferase activity of the MuRF1a (-638) reporter cotransfected with different *foxo* genes. Values are expressed relative to the luciferase activity of DMSO treated cells. ***, *p* < 0.001

**Table S1.**
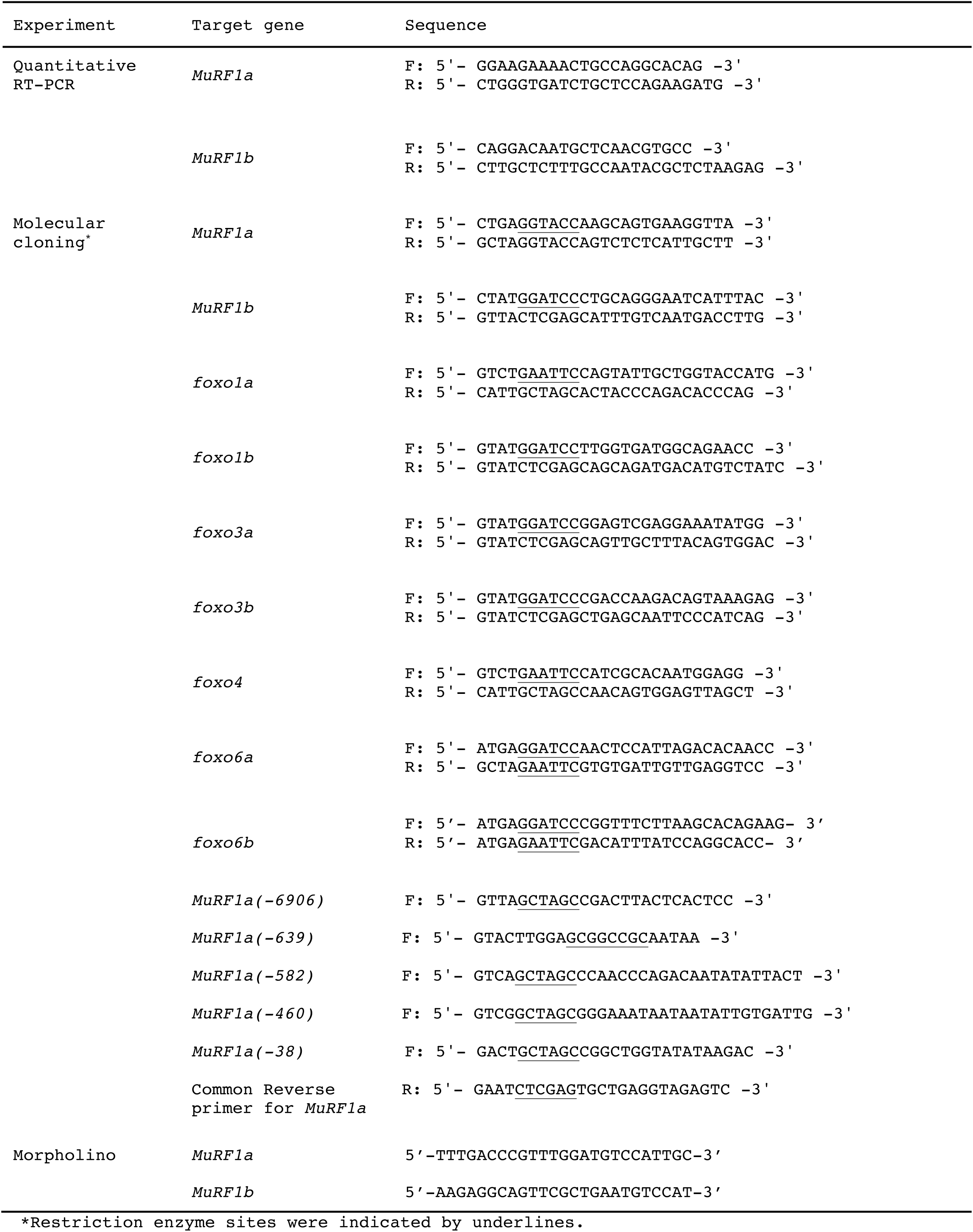
Primers and morpholinos used in this manuscript

